# Functional diversification enabled grassy biomes to fill global climate space

**DOI:** 10.1101/583625

**Authors:** Caroline E. R. Lehmann, Daniel M. Griffith, Kimberley J. Simpson, T. Michael Anderson, Sally Archibald, David J. Beerling, William J. Bond, Elsie Denton, Erika J. Edwards, Elisabeth J. Forrestel, David L. Fox, Damien Georges, William A. Hoffmann, Thomas Kluyver, Ladislav Mucina, Stephanie Pau, Jayashree Ratnam, Nicolas Salamin, Bianca Santini, Melinda D. Smith, Elizabeth L. Spriggs, Rebecca Westley, Christopher J. Still, Caroline A.E. Strömberg, Colin P. Osborne

## Abstract

Global change impacts on the Earth System are typically evaluated using biome classifications based on trees and forests. However, during the Cenozoic, many terrestrial biomes were transformed through the displacement of trees and shrubs by grasses. While grasses comprise 3% of vascular plant species, they are responsible for more than 25% of terrestrial photosynthesis. Critically, grass dominance alters ecosystem dynamics and function by introducing new ecological processes, especially surface fires and grazing. However, the large grassy component of many global biomes is often neglected in their descriptions, thereby ignoring these important ecosystem processes. Furthermore, the functional diversity of grasses in vegetation models is usually reduced to C_3_ and C_4_ photosynthetic plant functional types, omitting other relevant traits. Here, we compile available data to determine the global distribution of grassy vegetation and key traits related to grass dominance. Grassy biomes (where > 50% of the ground layer is covered by grasses) occupy almost every part of Earth’s vegetated climate space, characterising over 40% of the land surface. Major evolutionary lineages of grasses have specialised in different environments, but species from only three grass lineages occupy 88% of the land area of grassy vegetation, segregating along gradients of temperature, rainfall and fire. The environment occupied by each lineage is associated with unique plant trait combinations, including C_3_ and C_4_ photosynthesis, maximum plant height, and adaptations to fire and aridity. There is no single global climatic limit where C_4_ grasses replace C_3_ grasses. Instead this ecological transition varies biogeographically, with continental disjunctions arising through contrasting evolutionary histories.

**Significance statement:** Worldviews of vegetation generally focus on trees and forests but grasses characterize the ground layer over 40% of the Earth’s vegetated land surface. This omission is important because grasses transform surface-atmosphere exchanges, biodiversity and disturbance regimes. We looked beneath the trees to produce the first global map of grass-dominated biomes. Grassy biomes occur in virtually every climate on Earth. However, three lineages of grasses are much more successful than others, characterizing 88% of the land area of grassy biomes. Each of these grass lineages evolved ecological specializations related to aridity, freezing and fire. Recognizing the extent and causes of grass dominance beneath trees is important because grassy vegetation plays vital roles in the dynamics of our biosphere and human wellbeing.

## Introduction

The global distribution of terrestrial biomes determines global patterns of carbon storage and biodiversity (1). Delineation of biome distributions is crucial because it underpins evaluations of vegetation feedbacks on climate (2), extinction threats for biodiversity (3), and strategies for monitoring and reversing land-use change and degradation (4). Global studies of biome distributions typically focus on forests and trees (4-6), following the long-established paradigm in modern ecology of deterministic relationships between forest distributions and environment (7). Within this paradigm, there is a widely held perception that grassy vegetation only occupies semi-arid climates. However, it is increasingly recognized that biome limits are not deterministically linked to climate but arise from multi-directional feedbacks between plant functional traits, environment, and disturbance. These processes operate over evolutionary and ecological timescales (8) creating biogeographic contingencies in biome-environment relationships (9).

Grassy biomes require open-canopied tree layers (or no tree layer) to permit enough light to penetrate for grass photosynthesis. As a result, grasses dominate the ground layer when the rate of woody plant recruitment and growth is limited by climate, soil, drainage, disturbance conditions or light competition (10-12). “Grassy biomes” defined in this way include tropical savannas, montane grasslands, grassy deserts, temperate steppe grasslands, boreal parklands, and many temperate woodlands. The distinction of whether the ground layer is dominated by grasses (Poaceae) is fundamental to understanding global relationships among plants, climate, and disturbance (13). While, both trees and grasses are clearly important in driving vegetation dynamics, grass dominance causes a fundamental shift in disturbance regimes, whereby the consumption of ground layer biomass by fire and grazing reinforces grass dominance and maintains open tree canopies (10). Grass cover and biomass in the ground layer also affects surface energy, carbon, nutrient and water cycling by, for example, altering rates of decomposition, water infiltration and absorption of sunlight. Grass dominance therefore leads to novel ecological processes and properties in the Earth System, including frequent fire and grazing by mammals (14).

During the Cenozoic grasses displaced forests and shrublands by altering disturbance regimes at large scales across tropical and temperate regions (14, 15). The global expansion of grassy vegetation enabled major faunal and floral radiations (14, 16), and is linked to events in human behavioral evolution (17, 18). Today, natural grassy biomes provide grazing lands, water resources and numerous ecosystem services that directly support over a billion people (19). Yet, despite this social and economic significance, and the profound disturbance feedbacks engendered by grassy vegetation (20), understanding of grassy biomes is geographically biased towards few regions (e.g., South and East African savannas, North American grasslands), with the global limits of grassy biomes poorly defined.

When considering the limits to grassy biomes, the grass diversity present in a system is generally reduced to a distinction between species using the C_3_ or C_4_ photosynthetic pathways. If all else is equal, C_4_ grasses should outcompete C_3_ grasses under conditions of high light and temperature as well as low CO_2_ (21-23). This physiologically based model explains, in general terms, how C_4_ grasses dominate tropical regions and C_3_ grasses dominate temperate and high-altitude environments under current atmospheric CO_2_ levels [ppm ≈ 408]. The physiological mechanisms underpinning this model have critical impacts for predicting vegetation trajectories with global climate and atmospheric CO_2_ changes (22) yet attempts to parse the consequences of grass physiology for global vegetation is often reliant on sparsely validated modelling (24). Further, a focus on photosynthetic type belies the rich phylogenetic diversity within grasses independent of photosynthetic pathway (14). Grasses are unusual among vascular plants because C_4_ photosynthesis evolved in up to 24 independent lineages (25), conferring unique ecological characters to each C_4_ lineage inherited from its C_3_ ancestors (26). Photosynthetic type therefore interacts with different combinations of other functional traits to determine plant performance under varied environmental conditions (27, 28), but the influence of these interactions on the global biogeography of grassy biomes is unknown.

Here, we focus on grass-dominated systems to address three questions. First, what are the global limits of grassy biomes? Second, to what extent is grassy biome structure contingent on evolutionary history, whereby independent phylogenetic lineages characterize grassy biomes on each continent? Finally, how do functional traits of the descendant species of each lineage relate to climate and fire? Our findings have significant implications for the representation of terrestrial vegetation processes in Earth System Models.

### Identifying grassy biomes

Our dataset provides the first spatially explicit, functional classification of grassy vegetation at the global scale (Figs. 1 and S1). Necessarily, our approach that focusses on the ground layer contrasts with efforts to map biomes using remotely sensed tree cover or biomass (4, 29). Such studies generally misclassify extensive areas of tropical savanna as forest or degraded forest (30, 31). Global synthesis of grassy biomes has been prohibited as satellite remote sensing does not see through a tree canopy. Therefore, we mapped grassy formations by integrating and re-classifying 20 existing national and regional vegetation maps produced using botanical data and detailed vegetation descriptions (see Methods and SI).

**Figure 1:**
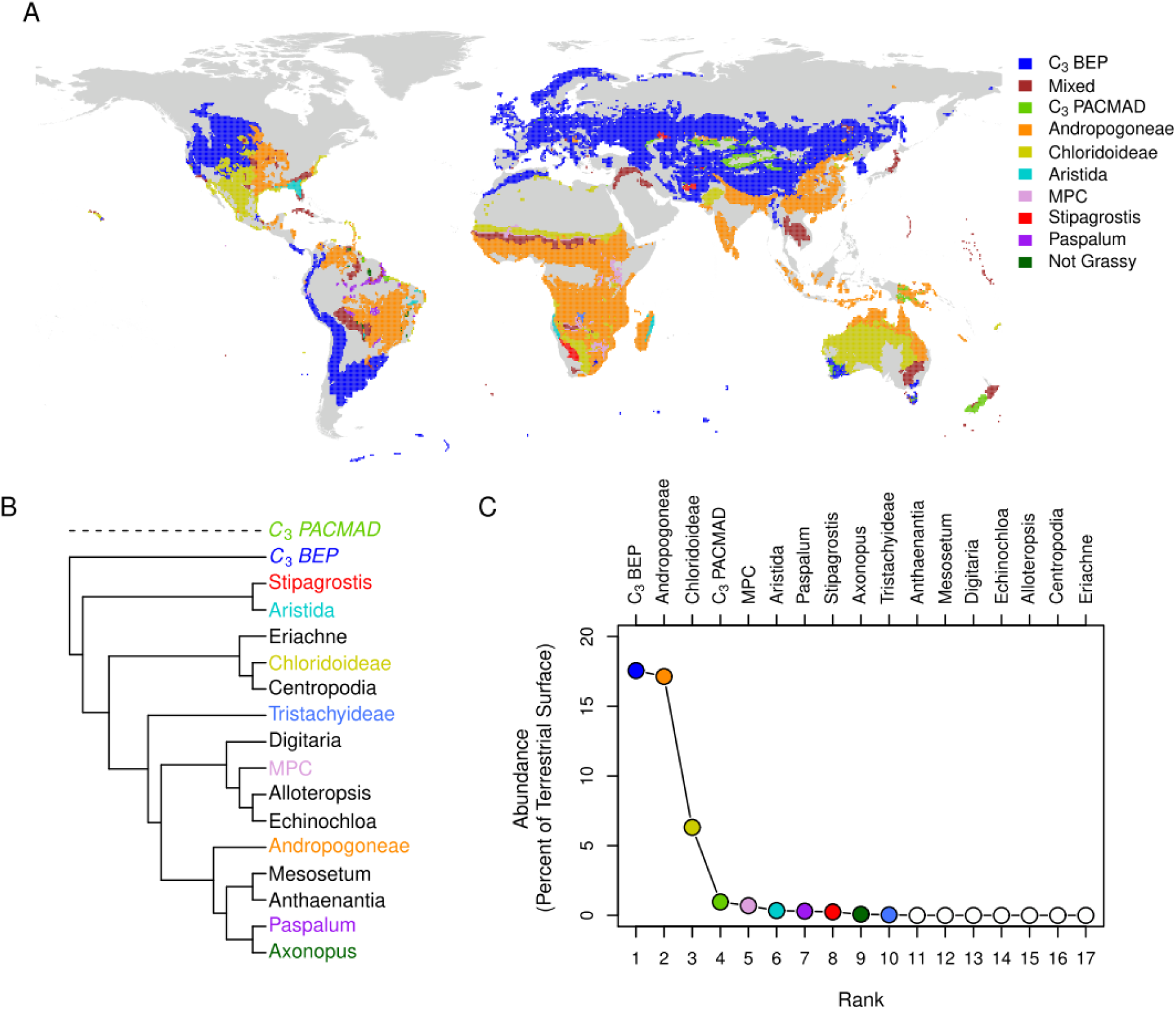
Global distributions of grassy biomes and dominant grass lineages. A. Grassy biomes coded by the C_4_ and C_3_ grass lineages dominating each vegetation formation. B. Relationships among the dominant C_4_ grass lineages, with colours matching those used on the map. The phylogeny is based on (25) and for simplicity excludes C_3_ PACMAD sister clades. C. Rank-abundance curve for C_4_ and C_3_ grass lineages at the global scale, ordered by the proportion of the terrestrial surface dominated by each lineage.

What is a grassy biome? We defined vegetation units as grassy where the ground layer is characterized by Poaceae and where grasses comprised > 50% of ground layer cover based on descriptions within vegetation maps and associated literature (see Methods and SI). A relatively small set of species often accounts for the majority of biomass in plant communities, whether these are communities dominated by trees or grasses, and these species exert major controls over ecosystem processes (32) and ecosystem services (33). Focusing on dominant and characteristic species provides one way to explore links between evolutionary history and ecosystem ecology at large scales (14). Through this process we identified 1,154 grass species (~10% of the total grass flora) characterizing grassy vegetation.

## Results and Discussion

### Global limits of grassy biomes

Grasses can dominate ground layer vegetation in all but the coldest and driest climates on Earth (Figs. 1 - 2). We estimate that vegetation with a grass-dominated ground layer originally covered ~ 41% of the vegetated land surface, although much is now under cultivation. Critically, grasses can dominate the ground layer in every climate where woody vegetation can persist (Figs. 1 - 2). While steppe grasslands and prairies occupy a large fraction of the global land area in dry temperate climates (Figs. 2, S2-S3), and savannas and grasslands occupy most of the tropics, grass-dominated ground layers occupy extensive areas in any other part of the vegetated global climate space (Fig. 2 and Fig. S4).

**Figure 2:**
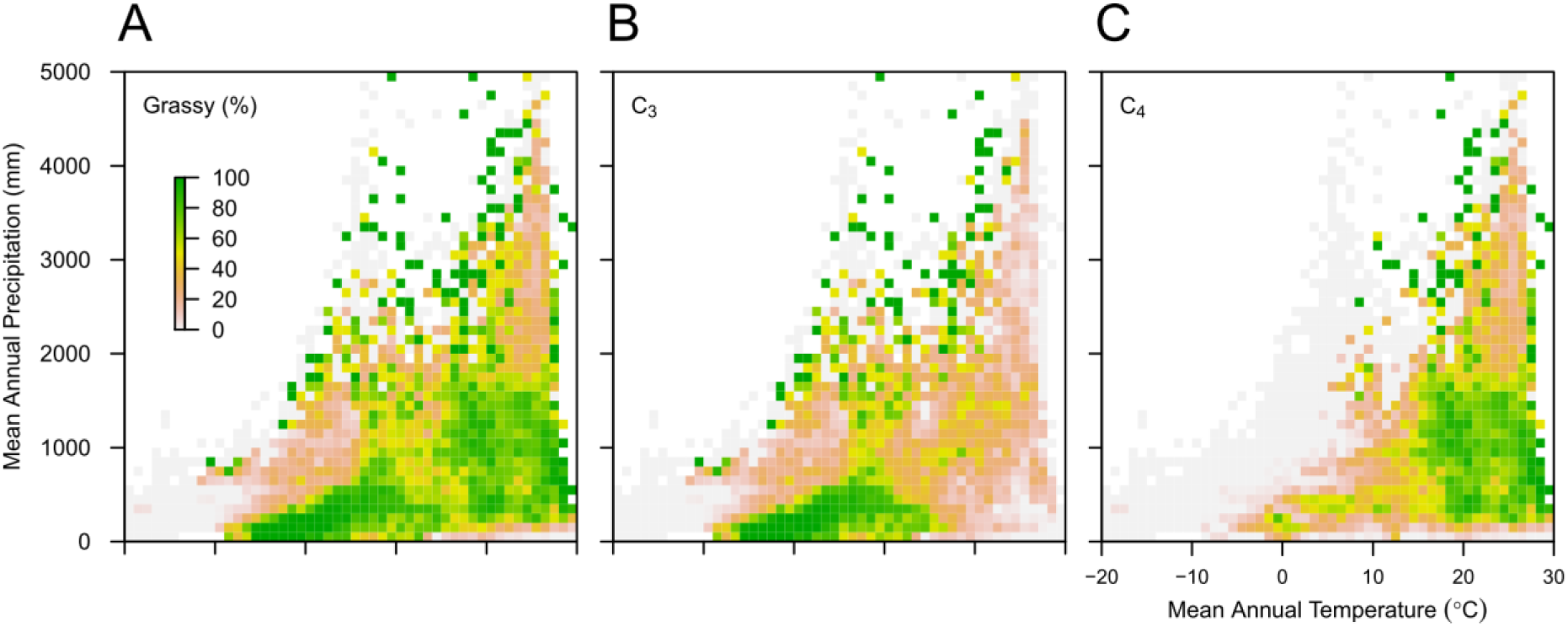
Grassy biomes in global climate space. Based on data at a 0.5-degree resolution, where data has been binned in 1^°^ mean annual temperature (MAT) and 100 mm mean annual precipitation (MAP) intervals. Colour ramp shows the relative proportion of the global climate space for that MAP × MAT bin occupied by grassy biomes. Grey shading represents the vegetated land area. Data shown here link to Figure S1 showing the total vegetated land area within each climate interval.

Members of 16 independently derived C_4_ grass lineages dominate within at least one vegetation unit worldwide (Fig. 1). However, two C_4_ lineages and one C_3_ lineage dominate over 88% of the land area of grassy vegetation: C_4_ Andropogoneae, 37% (1189 species in the lineage); C_4_ Chloridoideae, 14% (1601 species in the lineage); and C_3_ BEP, 38% (Fig. 1). The vast majority of C_3_ BEP taxa belong to Pooideae (4234 species in the lineage). In contrast, C_3_ species of the PACMAD clade dominate only 2% of grassy biomes (Fig. 1); these are the closest relatives of C_4_ grasses and are restricted to warm, wet areas (Figs. S2-S4). Of the remaining area of grassy vegetation, 6.6% is characterised by a mix of lineages, and the rest dominated by 13 other, independently derived, C_4_ lineages (Fig. 1 and Table S1).

The three dominant lineages sort in climate space. C_3_ Pooideae dominate cooler, drier climates, whereas C_4_ Andropogoneae and Chloridoideae dominate grassy biomes in warmer climates (Figs. 3, S3-S6). However, precipitation sorts the C_4_ lineages, with peak dominance of Andropogoneae occurring at ~ 1200mm MAP (Figs. 4, S3-S5), coinciding almost precisely with the global peak in fire frequency (Fig. 4). This is also the climate space where disturbance-driven feedbacks are considered to play a major role in maintaining open (i.e., grassy) or closed (i.e., woody) vegetation (34). In contrast, the peak dominance of Chloridoideae occurs at ~350mm MAP (Figs. 4, S3-S4), within semi-arid climate zones occupied by both dry savannas and shrublands/thickets e.g., (35). Temperature seasonality also differs among the C_4_ lineages, with Chloridoideae dominating in regions with strong seasonality, and Andropogoneae dominating in more aseasonal environments (Fig. S5).

**Fig. 3.**
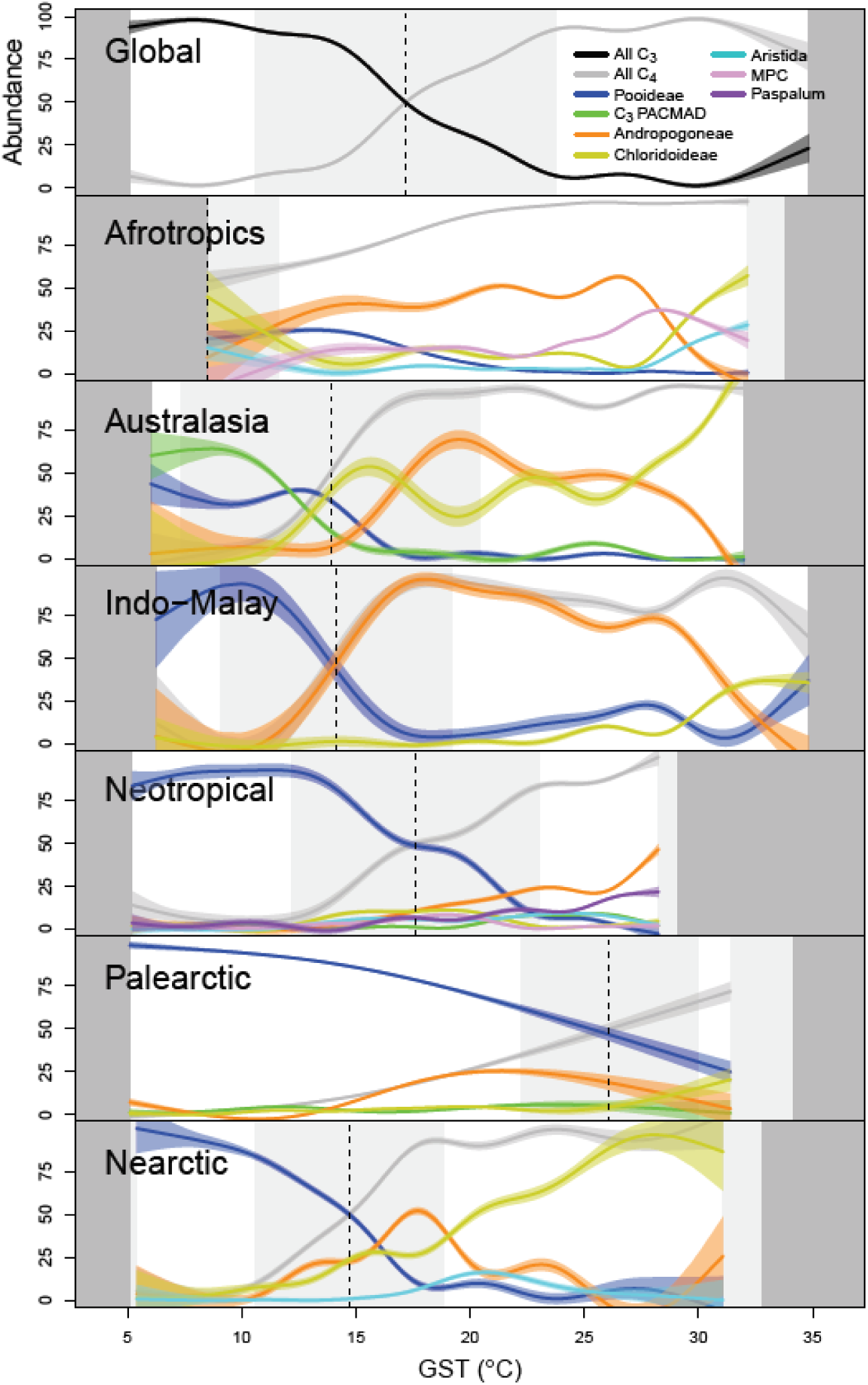
Continental disjunctions in lineage-growing season temperature relationships. The distribution of grass lineages relative to growing season temperature (GST) in degrees Celsius globally (top panel) and then showing the variation in estimated C_3_-C_4_ crossover temperatures by geographic realm. Distributions was fitted using generalized additive models and the crossover temperature calculated as the point where modelled C_4_ grass abundance is 50%, based on the mapping in Figure 1. The fitted lines and confidence intervals are shown in different colours for each lineage, with the legend on the figure.

**Figure 4.**
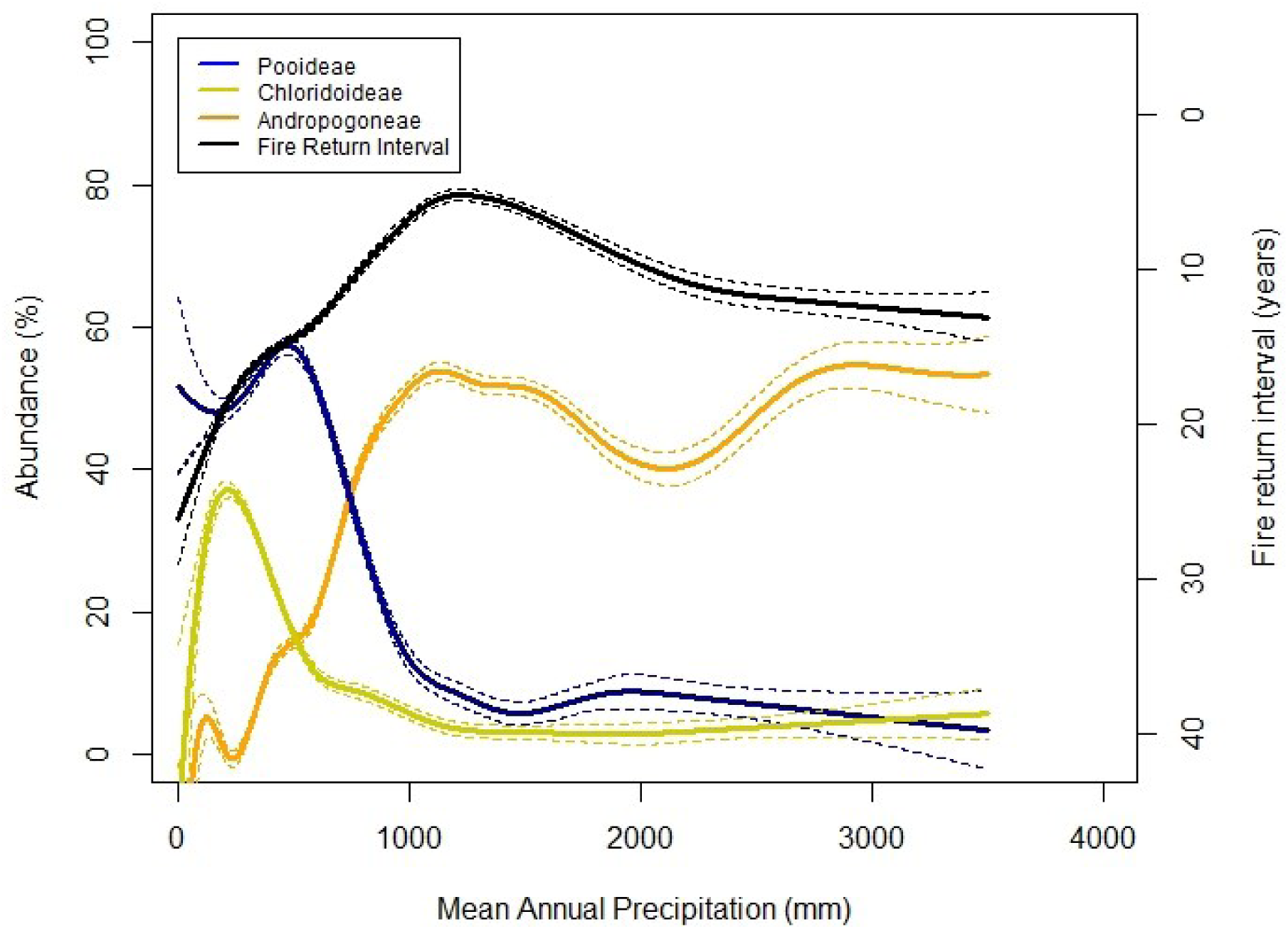
Global relationships between rainfall, fire and grass lineages. The global relationships of grass lineage abundance relative to MAP using a generalised additive model, showing 95% confidence intervals. The right-hand axis is the global relationship between fire return interval and MAP for grassy biomes and is inverted to reflect the inverse relationship with MAP. The global peak in fire activity coincides with the global peak in dominance of Andropogoneae grassy biomes. In contrast longer fire return times are associated with dominance by Pooideae and Chloridoideae.

### Continental disjunctions in C_3_ and _4_ lineage distributions

Globally, the mean growing season temperature where dominance of grasses transitions from C_3_ to C_4_ types varies starkly among continents, from 8.5-26.1 °C, with a global mean of 17.2 °C (Fig. 3). Lineages using C_3_ and C_4_ photosynthetic pathways are clearly sorted by growing season temperature and mean annual temperature (Figs. 3 and S3-6). The C_3_ Pooideae lineage has specialized and radiated in cold environments by evolving physiological cold acclimation to protect tissues from freezing damage, and vernalization to synchronize flowering with the growing season (36, 37). Conversely, in tropical regions, the repeated evolution of C_4_ photosynthesis appears vital in expanding the range of grassy biomes, by enabling colonization of hot, high light, and seasonally dry habitats across a wide span of rainfall (38, 39).

The C_3_ Pooideae occupy regions with lower winter temperatures and shorter droughts than the C_4_ lineages (Fig. S7). C_3_ Pooideae dominate grassy biomes to much higher temperatures in the Palearctic than the Nearctic realm, although distributions of C_4_ Andropogoneae and Chloridoideae in these realms are similar (Fig. 3). Conversely, C_3_ Pooideae are confined to the geographically restricted colder parts of the Afrotropics and Indo-Malay realms, and C_4_ Andropogoneae dominate at much lower temperatures in these regions (Fig. 3). The sorting of C_3_ and C_4_ grass species along local and regional temperature gradients is well established (40, 41), and the crossover temperature can be modified by ecosystem factors (e.g., tree cover) (42). However, our observations are broadly consistent with model predictions of carbon assimilation (22, 23, 43), as modeled crossover temperatures under low light conditions and modern CO_2_ levels occurs at ~20-22 °C.

In our data, some species of both Andropogoneae and Chloridoideae lineages have adapted to low mean annual temperatures and may persist in grassy vegetation within cool parts of each realm (e.g. Fig. 3). Given equal investment in the carbon-fixing enzyme Rubisco, a relatively low canopy leaf area and sunny conditions, a C_4_ canopy can theoretically achieve higher total daily rates of photosynthesis than a C_3_ at any temperature (37). In this case, the primary limitation on canopy carbon uptake becomes light-mediated damage during low temperature extremes (44), although C_4_ photosynthesis is energetically expensive. Low temperature tolerance may be absent from most C_4_ species as C_4_ photosynthesis evolved in the tropics (38).

### Trait combinations of each lineage

Chloridoideae are distinguished from Andropogoneae in their occupation of regions with lower precipitation, higher daily variation in temperatures and longer droughts (Fig. S7). Further, these lineages are differentially associated with fire where Andropogoneae has the shortest fire return interval of 2 years, the peak occurrence of Chloridoideae is at an interval of 8 years, while in Pooideae the modal fire return interval exceeds 20 years (Fig. S7). Maximum plant heights of each lineage sort similarly, with values peaking at 1.5 m for Andropogoneae and 0.6 m for both Chloridoideae and Pooideae (Fig. S7). However, annual versus perennial life history is not globally relevant. The only significant areas dominated by annual grasses occurring at the margins of the Sahara Desert and West Africa, regions commonly considered as over-grazed.

80% of burned area globally occurs in the regions we see dominated by Andropogoneae (20) and differs from other C_4_ grass lineages with its greater average height and consequent rapid growth rates. Where rainfall exceeds 800 mm MAP in the tropics, soils are typically leached and infertile (45). Andropogoneae produce leaves with relatively high C:N ratios (46, 47), which resist rapid decomposition. The tall, erect architecture of these grasses produces a flammable well-connected fuelbed (48) and productive tropical environments, with an annual dry season of > 5 months (13), are primed to burn as the grass layer senesces. Experimental manipulations demonstrate that fire promotes dominance by Andropogoneae (46) and we see this mirrored at a global scale. Grass persistence in these competitive environments relies on the annual production of a new canopy and, in the absence of woody investment, dead biomass must either rapidly decompose, burn or be consumed by herbivores to avoid self-shading (11, 49). Andropogoneae are known to have morphological adaptations enabling tolerances and persistence to fire that are not commonly present in other grass lineages (49). Fire and other forms of repeated disturbance, such as grazing, are therefore crucial for grass-dominated systems to persist in high rainfall environments. While Andropogoneae appears to be the C_4_ lineage most closely associated with disturbance by fire, multiple lineages in the semi-arid African tropics appear linked to grazing tolerance (Fig S8, (50, 51)), and this may be due to the strength and form of environmental filtering associated with fire versus grazing, as well as the antiquity and biogeography of grazing pressure relative to fire.

### Implications

The Andropogoneae, Chloridoideae and Pooideae grass lineages dominate globally, via mechanisms encompassing plant production and competition, resilience to drought, freezing and disturbance. Why do three of the most diverse grass lineages characterise grassy biomes? Does diversity beget ecological success or does success beget diversity? Early diversification may have enabled ecological success, such that ecological speciation allowed each lineage to radiate across broad environmental envelopes (an ecological mechanism). Alternatively, a neutral mechanism of a long history of diversification may have led to high diversity as Andropogoneae and Chloridoideae are the oldest C_4_ lineages. Across our dataset, evidence for this is equivocal. We list 8.8% of all grass species and within lineages: Andropogoneae, 14.5%; Chloridoideae, 6.5%; Pooideae, 10.8%. Perhaps ecological success facilitated diversification, such that large geographical ranges enabled by unique adaptations made the isolation of populations and allopatric speciation more likely (a geographic mechanism). The rapid spread of the cosmopolitan *Themeda triandra* from Asia to Africa in < 500,000 years supports this idea (52). Resolving the relative role of these mechanisms requires comparative phylogenetic analyses of the relationships among ecology, functional traits, range sizes and diversification rates.

The biogeographic contingencies described here in crossover temperatures align with emerging evidence that regional evolutionary and environmental histories have been important modifiers of biome-climate relationships (9, 53). However, the rapid rates of dispersal observed in grasses (52), along with their short generation times (49), raises critical questions about whether the biogeographic contingencies observed in woody plants should be mirrored in grassy communities.

Global change will rapidly modify the existing global distribution of grassy biomes. First, environmental change can alter feedbacks between grasses and woody plants via changes in the processes limiting the growth and mortality of woody plants. For example, rising CO_2_ is hypothesised to increase tree recruitment in savannas and forest margins (54, 55), while extreme drought events and warming may cause forest dieback on large scales (56), where each process has feedbacks with fire leading to ongoing biome shifts (57). Second, environmental changes will shift the community composition of grass communities. Our analysis points to globally important ecotones between C_3_ and C_4_ likely to be influenced by rising CO_2_ and temperature (58), but these are better conceptualised as the boundary between Pooideae and Chloridoideae in arid and semi-arid regions or regions of high grazing pressure, and Pooideae and Andropogoneae in wetter regions. An experimental CO_2_ manipulation in dry mixed prairie found elevated CO_2_ favoured a Pooideae dominant over a Chloridoideae dominant, with rising temperature having the opposing effect (59). Conversely, in a mesic tallgrass prairie, an Andropogoneae dominant displaced a Pooideae dominant in competition under elevated CO_2_ via improved water relations (60). In each case, C_4_ photosynthesis was one trait among many that influenced dynamic environmental responses. Finally, the boundary between Andropogoneae and Chloridoideae is more likely to be influenced by changes in rainfall amount and seasonality, along with shifting fire and grazing regimes that can be directly altered by people at small and large scales.

## Conclusions

The previous lack of synthesis in biome limits between grasses and woody plants constrains our understanding of how ecological and evolutionary processes determine the sensitivity of vegetation to global change. We have shown that divergent evolutionary histories and unique functional trait combinations have enabled three major grass lineages to dominate grassy biomes across global climate space. Local dominance by each lineage brings differing sensitivities to alternative global change drivers.

## Acknowledgements

This research is a product of the National Evolutionary Synthesis Center (NESCent) working group led by CPO, CAES, and CJS. DG was supported by a NESCent graduate fellowship and NSF award 1342703. CPO was supported during the preparation of this manuscript by Natural Environment Research Council grant (NE/I014322/1). Zhiyao Tang helped to obtain the China map. Nikolai Ermakov and Daoud Rafikpoor provided shape files of mapping data for Russia and Afghanistan, respectively. Anita Smyth assisted in obtaining the Aekos data. Les Powrie, and Mike Rutherford assisted in obtaining the plot data from South Africa. The South African National Biodiversity Institute and the South African Biodiversity Facility are thanked for the use of data/information supplied by SANBI from digitized collections. This work forms part of the “The National Vegetation Map” coordinated by the South African National Biodiversity Institute.

## Author Contributions

CERL, DMG, KJS, TMA, WB, ED, EJF, WH, LM, SP, JR, BS, MS, ES, RW, and CPO compiled the data. CERL, DMG, KS, DG and TK analysed the data. SA contributed fire data. CERL and CPO designed the study and wrote the paper with text contributions from DMG and KJS. All authors contributed comments on a draft of the paper. DMG and CERL perfected the figures.

## Methods

### Classifying grassy biomes

Data from 20 vegetation maps derived from botanical information, or a combination of botanical and geographic information, were integrated to delineate grassy biomes (references for these maps are listed in the Supplementary Information). The result was a global map of grassy biomes resolved into 1,635 discrete vegetation units, each defined by its characteristic grass species, which formed a list of 1,154 species (accounting for synonymy) found commonly across global grassy biomes.

Vegetation maps are generally based on botanical survey and geographic analysis, combined with expert input, that cluster species composition and vegetation structure to define unique vegetation units. We compiled the ground layer information for the vegetation units in each map to identify the grass species considered to characterize a vegetation unit. To determine whether vegetation units were naturally dominated by grasses, we developed a set of criteria. First, artificial vegetation units were defined as those plowed or sown for agriculture and where humans are planting species that would not otherwise occur. We retained data for this analysis of only natural formations. Second, based on the vegetation descriptions we determined whether > 50% of the relative ground cover or biomass was derived from grasses. We used this definition in place of ‘Is there a continuous grassy ground layer?’ because low herbaceous cover in predominantly grassy vegetation would present a problem with the classification of desertic and arid environments. Vegetation units were considered grassy deserts where the total above-ground biomass was considered <50 g m^2^, or where total ground cover <25%, throughout the year. Finally, we retained all formations where grasses were the dominant component of the ground layer, irrespective of tree cover. Numerous grassy biomes, such as tropical savannas and woodlands, may be characterised by up to 80% tree cover, but behave functionally as savannas due to a contiguous grassy ground layer (13, 35). Where necessary, we sourced additional information from published vegetation descriptions and analyses to attribute key grass species to a grassy vegetation unit. Additionally, vegetation units could be classified as mosaics with patches of closed canopy vegetation intermingled with open vegetation, e.g. across the Steppe region of Russia.

### Mapping grassy biomes

The vegetation maps we used as sources were developed throughout the 20^th^ century. While this method provides an incomplete global coverage, we integrated available state-, country- and continent-level mapping to assemble what we consider to be the most robust map possible of the limits of grassy vegetation, where both vegetation characteristics and key constituent species could be identified. We were obliged to use the WWF Ecoregions map (61) where no other mapping was available. We re-assessed this global map to re-define units as grassy or not based on the criteria outlined above.

To quantify the global limits of grassy vegetation according to grass lineage, we gridded the mapped data compilation at 0.5 degrees resolution. We calculated the proportion of each 0.5-degree grid cell occupied by grassy polygons. Using the grass phylogenetic and trait information compiled, we then calculated the occupancy of grassy polygons by photosynthetic type, annual/perennial life history, grass lineage, and mean maximum grass height. These data are not the same as a classic concept of abundance or dominance but are a relative measure of the likelihood of occupancy measured from zero to unity. We undertook a validation of our map compilation described in the Supplementary Information and in Figure S9.

### Phylogenetic and plant trait information

We cross-referenced our species list to a taxonomy of accepted scientific names (GrassBase, http://www.kew.org/data/grasses-syn/cite.htm) and a recent accepted phylogeny from the Grass Phylogeny Working Group (25) to eliminate synonymy and link species to descriptions of evolutionary history and functional traits. Functional traits considered were: C_3_/C_4_ photosynthetic pathway, maximum plant height, annual/perennial life history, and tolerance of climatic extremes and fire frequency. C_3_ species were divided amongst two groups: a polyphyletic group belonging to the PACMAD clade (including the C_3_ sister groups for all C_4_ lineages); and the monophyletic BEP clade, a C_3_ outgroup to PACMAD, including bamboos, rice relatives and Pooideae species. C_4_ grass species were attributed to one of 24 independently evolved grass lineages. Maximum plant size is a major axis of plant trait variation at a global scale (62), with maximum culm height in herbaceous grasses reflecting annual rates of height growth, as most grasses annually senesce their canopy (49). Height also describes differences in life history strategies related to light competition and flammability and grazing tolerance (49). We included annual/perennial as while most grasses reach sexual maturity in < 1 growing season, perennial grasses can be long-lived. Plant longevity is an effective strategy for occupying space in competitive environments (63). We summarized these data for each grassy vegetation unit based on the grass species listed as characteristic of each unit.

For the Poaceae species that we listed, we extracted all available georeferenced occurrence records from the Global Biodiversity Information Facility (GBIF) web portal (http://www.gbif.org/; accessed January 2014) and cleaned these data to ensure longitude and latitude values were viable and to two decimal places. Species distributions were standardised against descriptions of distributions in Grassbase using TDWG regions. For this subset of species produced via distribution records, median fire return intervals were calculated at a species level following the methods of Archibald *et al.* 2010 (64). Information on fire date was extracted for each GBIF location from the MODIS global monthly burnt area (MCD45A1) satellite data product. To calculate climatic extremes for these same species, the WorldClim dataset (www.worldclim.org) was used to obtain species median values of minimum temperature (BIO6 variable) and seasonal drought length (calculated as the number of successive months where mean annual precipitation was below 30mm). These species level data were used to construct frequency histograms to examine lineage level variation in fire regimes and climate extremes (Fig S7).

### Environmental data used in global analyses

Our analysis aimed to elucidate lineage, climate and disturbance relationships, and whether biogeography impacts the C_3_-C_4_ crossover temperature. We used the WorldClim dataset at a 0.5 degree resolution to match the vegetation map, and extracted mean annual precipitation (MAP), rainfall seasonality, mean annual temperature (MAT) and temperature seasonality (www.worldclim.org). We used a rainfall concentration index to describe rainfall seasonality based on (35). Growing season temperature (GST) was calculated for each grid cell to quantify regional and global C_3_-C_4_ crossover temperatures. GST was calculated as the mean temperature across months with a greater than or equal to 5 degree mean temperature and at least 25 mm rainfall, and was calculated using WoldClim monthly climate normals (65).

A median fire return interval (FRI) is the number of years between fire events that represents the time period available for plants to grow. We used fire interval data from the 16 year MODIS fire datasets to fit Weibull distributions to 0.5° gridded data for the globe by using the method outlined in Archibald *et al.* 2010 (64). Tropical grasslands and savannas have the world’s shortest fire return times, due to rapid rates of fuel accumulation and a climate that supports frequent fire (annual dry seasons, warm climate and reliably seasonal rainfall) (20). Our dataset of estimated fire return times, while spatially biased, is therefore robust for grassy biomes.

Globally consistent data on present or past herbivore pressures are simply not available. We were obliged to restrict our analyses to Africa where efforts have been made to map mammalian herbivore pressures of both wildlife and livestock (66). We combined herbivore and fire data to assess links between lineage composition and disturbance. Soils data are not of sufficient quality to be meaningfully incorporated in analyses of this scale, despite being known to mediate local scale vegetation patterns (35). In our global analyses, we excluded grassy vegetation units defined as flooded, saline or edaphic, where the limits of these units are generally decoupled from climate.

### Analyses

First, we mapped the distribution of grassy biomes in geographic space according to lineage and photosynthetic type to calculate the land area occupied by different grass lineages in a “rank-abundance” style (Fig. 1). Grassy biome distributions were aligned with MAT (in 1°C intervals) and MAP (in 100 mm intervals) to construct “Whittaker” style plots of the limits of grassy biomes and of C_3_ and C_4_ photosynthetic types (Figure 2). These data were further decomposed to represent the climate space of 17 grass lineages including Pooideae that dominate grasslands (Figure S4). Data were also analysed by climate intervals of MAT and MAP to calculate the proportion of grassy land area occupied by each grass lineage within each climate interval, to consider the potential for deterministic links between climate and biomes (Figs. S3-S4).

Generalised additive models relating the distribution of lineages to growing season temperature, MAT and MAP across continents were fitted using the mgcv R package and the function predict.gam (67). Crossover temperatures plus standard deviations were calculated based on the temperature at which the predicted abundance of C_4_ dominance reached 50%. Random forest regressions (https://cran.r-project.org/web/packages/randomForest/randomForest.pdf) were used to examine the climate niche of key grass lineages and to infer correlations between four key climate predictors (MAP, MAT, temperature seasonality and rainfall seasonality). Models were constructed for six groups of interest: the C_3_ BEP, PACMAD and Pooideae lineages; and the independent C_4_ lineages Andropogoneae, Chloridoideae and MPC (Melinidinae + Panicinae + Cenchrinae) (25). Model fit was checked via a mean of squared residual test. The relative importance of each environmental correlate was computed with a mean decrease of accuracy test. The computed coverage response plots for each grass group was an adaptation of the evaluation strip method developed by (68). These plots demonstrate the non-linear relationships between environmental gradients and the various grass lineages. To produce these plots, an environmental dataset was simulated where the focal environmental variable is varied over its full environmental range and where, for each interval, the observed median of each of other environmental variables (median over areas where the focus environmental variable is within the interval) is returned. The displayed curve in each case is the prediction of our Regression Random Forest model over this simulated dataset. The process used bi-variate response curves, where two variables rather than one vary simultaneously. The 90^th^ quantile of a kernel density function (function kde2d from the R package ade4) was used to plot limits of grass lineages relative to herbivore abundance and fire frequency.

## Supplementary Figures, Tables and Information

### Development and validation of map compilation

Rarely, if ever, has this rich body of vegetation mapping research been integrated with Earth system science or evolutionary studies. This is perhaps because vegetation mapping is considered a descriptive natural science in an age of big data. The contiguous land mass covering the countries of China, Mongolia, the former Soviet Union, Afghanistan, Turkey and Europe are represented by detailed botanical data. The regions of Africa, North America, Mexico, Panama, Venezuela, Brazil, Argentina, Papua New Guinea, Indonesia, northern and western Australia are also well documented by botanical data. However, there is a general paucity of adequate vegetation mapping available across India, South-East Asia (Burma, Thailand, Laos, and Vietnam), Central America, and parts of South America (Chile, Peru, Bolivia, Uruguay, Paraguay, Ecuador, and Columbia). It is worth noting that given anticipated impacts of global change on the distribution and dynamics of vegetation, an absence of publicly available vegetation mapping for key regions such as South East Asia and the Andes should be of concern to many.

We undertook a validation process between plot data describing *in situ* grass abundance and our global species list. Using publicly available data that intersected with vegetation unit descriptions we found that, at the level of independent evolutionary lineages of grasses (i.e., subfamily), we had strong confidence in the geographic and environmental relationships we elucidate here (Fig. S9). To validate the classification of common grass species across regions, we compared the species list in each vegetation unit to a plot level database developed for validation purposes (Fig. S9). Plot data were sourced from the literature and vegetation databases and assembled by the authors (see references in the Supplementary Information). 110 vegetation units contained enough plot level data for validation analyses. To determine what taxonomic levels agree with plot data, the comparison was conducted at the species and subfamily levels. We also examined the agreement of our map and plot datasets at a functional level by comparing the attribution of photosynthetic type. From the 507 common grass species across these vegetation units, 88% of these species were present in the plot dataset of those appropriate vegetation types. This is a very high degree of overlap in species in our mapping classifications and plot data, especially considering the difference in scale between local species plots and large vegetation units. Furthermore, we found that vegetation types generally had similar percentages of characteristic grass species represented in their plot datasets, although the agreement was worse for particularly large and broadscale vegetation units. To validate the higher taxonomic classifications and plant functional type classifications of our map units, we compared the proportion each classification in plots (weighted by abundance) to the proportion of that classification in our map. Because these data are on the interval (0,1) we used beta regression to model this relationship. Beta regression can be interpreted much like logistic regression, except that it allows continuous values in the dependent variable. Proportions of Poaceae subfamilies and functional types showed that plot values were strongly predictive of classified values in our map (Fig. S9).

**Figure S1:**
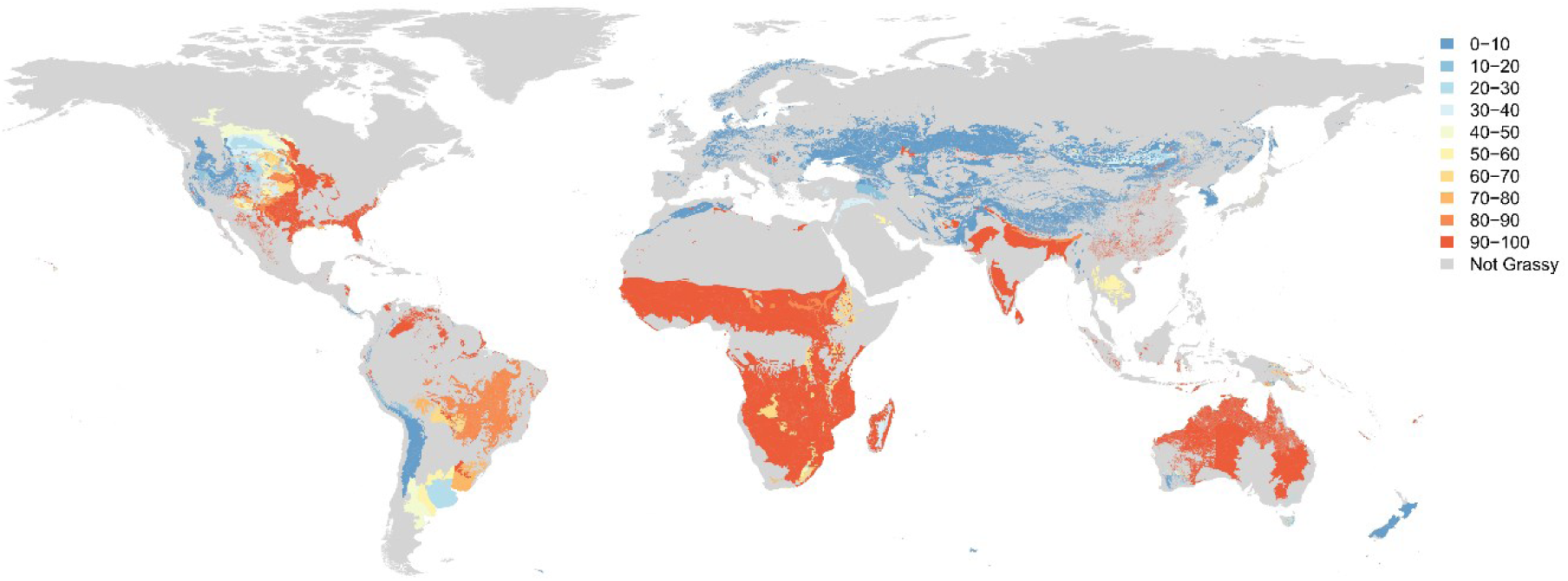
Global distribution of grassy biomes. The global map was derived as a composite from national and regional maps of vegetation that was gap-filled using the Ecoregions map (see Methods in the main text and references for all maps at the end of the Supplementary Information). Coloured areas show the extent of grassy biomes globally and dominance of these by C_3_ grasses and C_4_ grasses mapped at the scale of identified vegetation units (i.e., polygons). Red = High proportion of C4 grasses. Blue = High proportion C3 grasses. Datum: WGS84.

**Figure S2.**
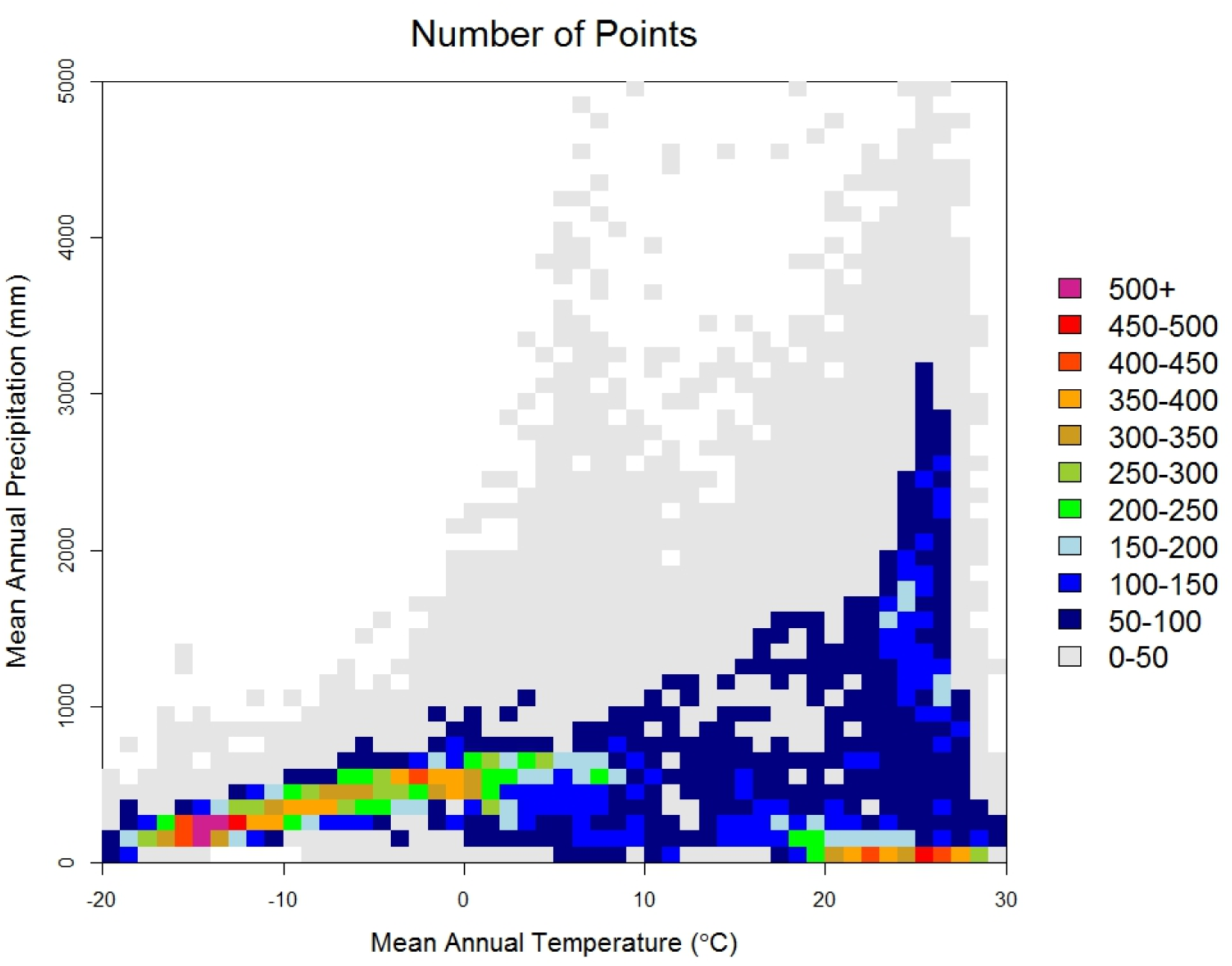
Global vegetated land area as related to Mean annual temperature and mean annual precipitation. Mean annual precipitation is in 100 mm bins, while temperature is in 1oC bins. The color ramp represents the number of 0.5 degree points in 100mm x 1oc unit of climate space. Note the grey background that highlights the global extent of climate space where these temperature – precipitation combinations are essentially rare on the vegetated land surface. The color ramp from dark blue to purple represents an increasing density of points in a given climate bin.

**Figure S3:**
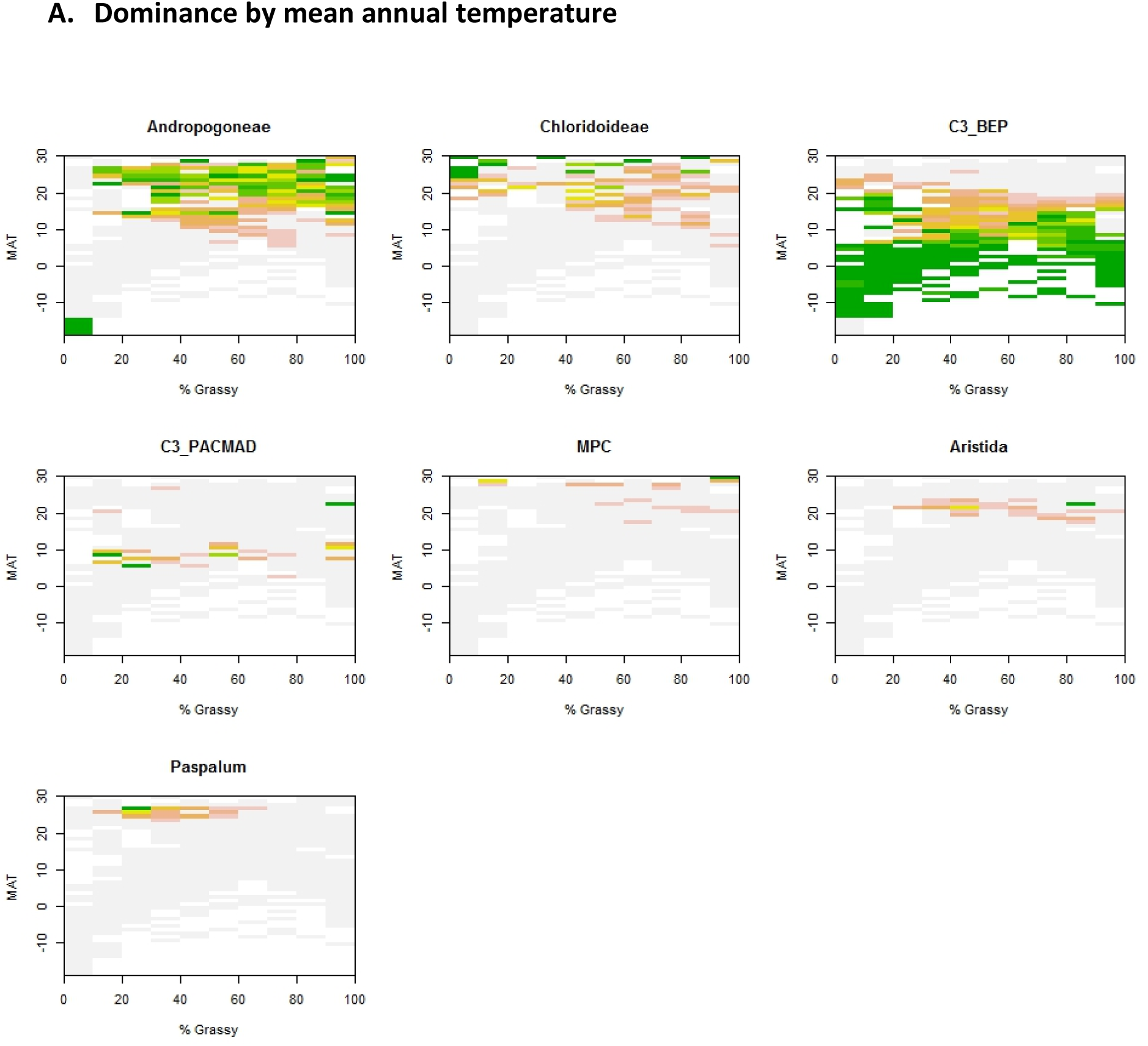
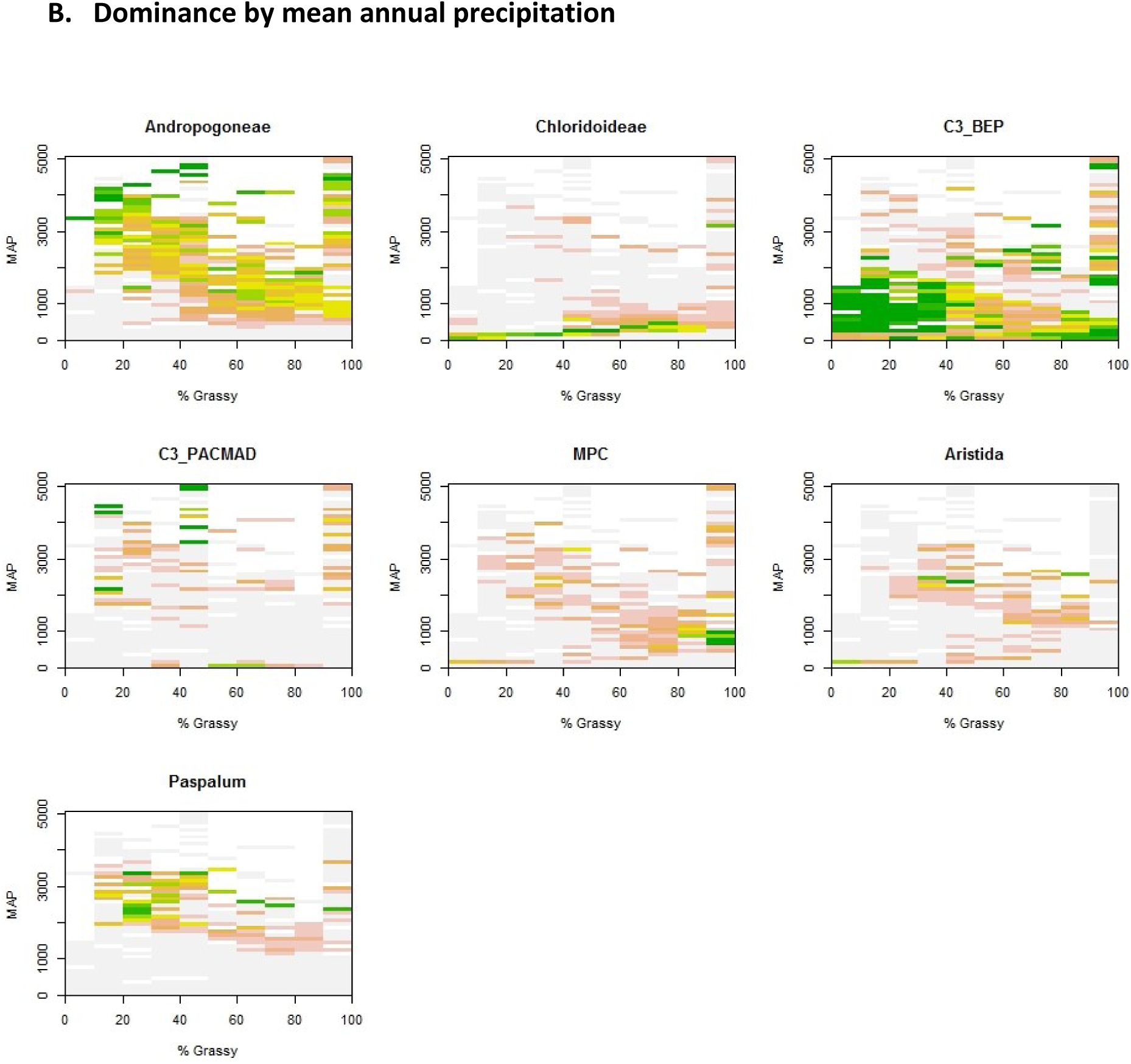
Global abundance of the main grass lineages by temperature and rainfall. Colour scale indicates the proportion of 0.5° grid squares dominated by each lineage at the global scale for each of (A) mean annual temperature and (B) Mean annual precipitation. These plots demonstrate that, in cool, dry regions where the C3 Pooideae lineage is concentrated, it tends to be the only grass lineage present, and this lineage dominates that climate space. These can be considered as deterministic grasslands. In contrast, the heterogeneity of the dominance of C_4_ Andropogoneae and C_4_ Chloridoideae lineages across climate space could suggest that the grassy biomes where these lineages are found are not deterministic, and dominance may be driven by processes other than climate.

**Figure S4:**
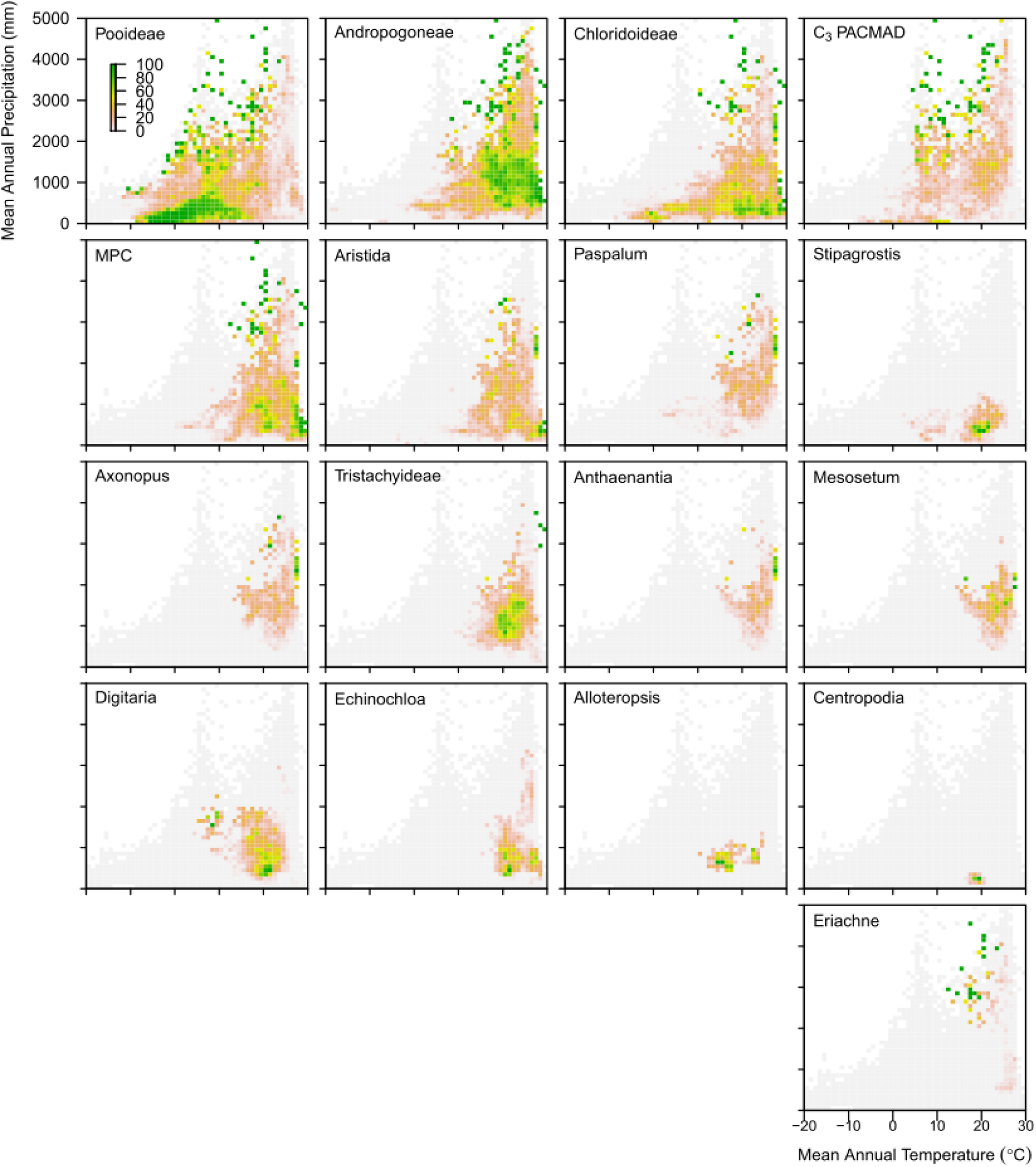
Concentration of 17 grass lineages in climate space. This figure builds on S1 – S2 by again highlighting the climate space characterised by different grass lineages. It is very clear that C3 PACMAD dominance is highly restricted to warmer wetter parts of climate and we know from S1 that geographically these combinations of temperature and precipitation are limited. These figures also again highlight the wide distribution of Pooideae, Andropogoneae and Chloridoideae.

**Figure S5:**
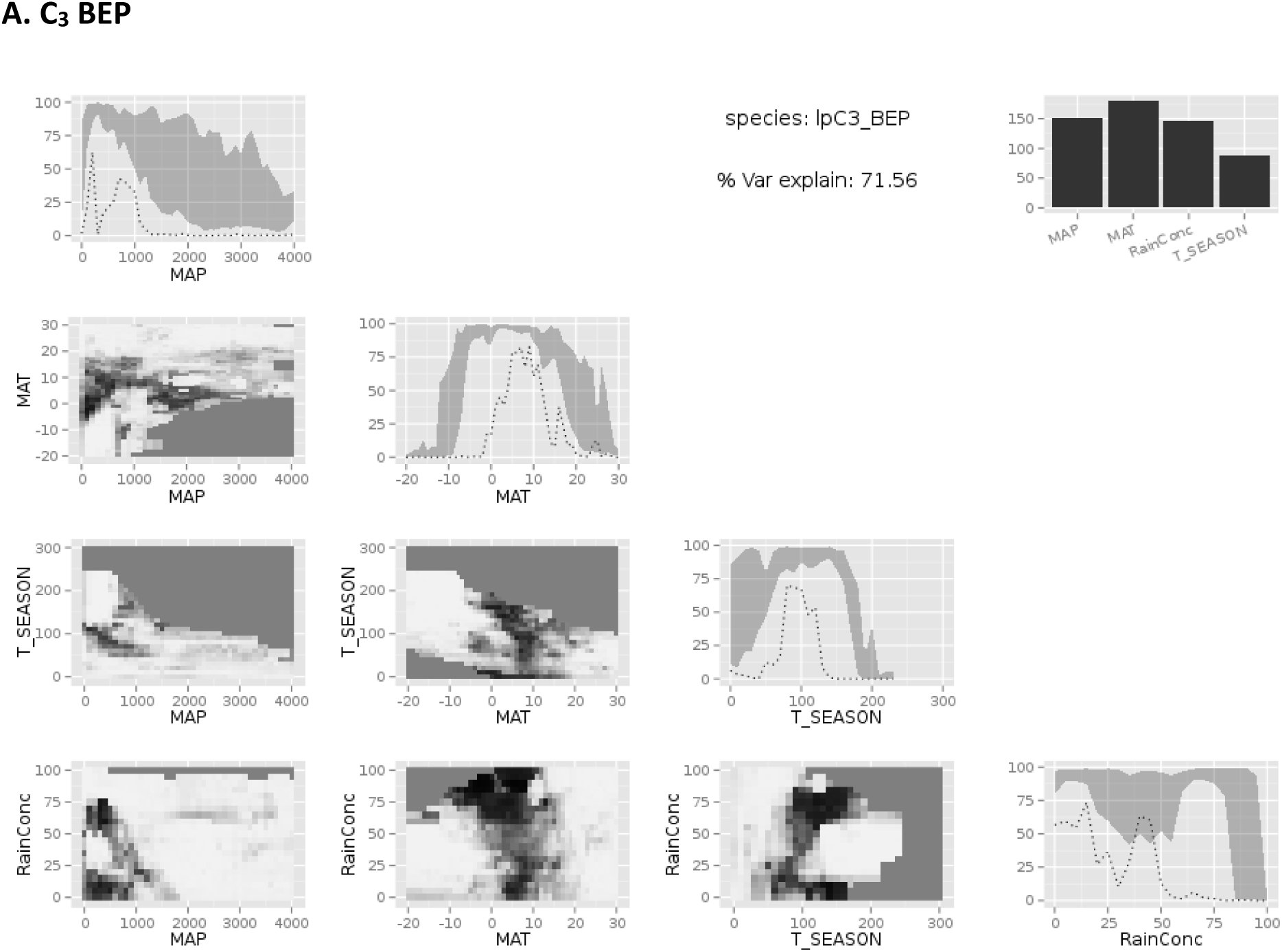
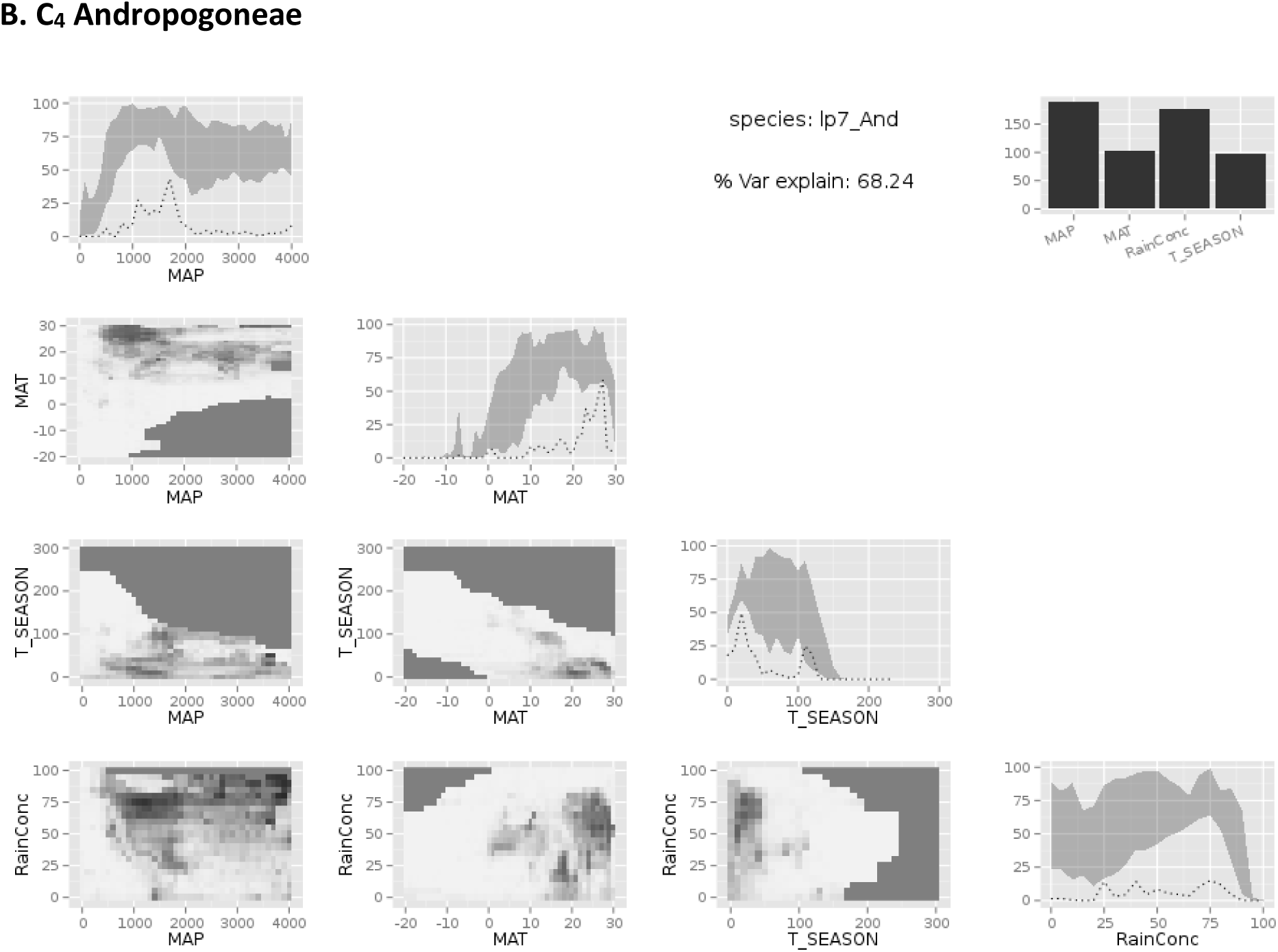
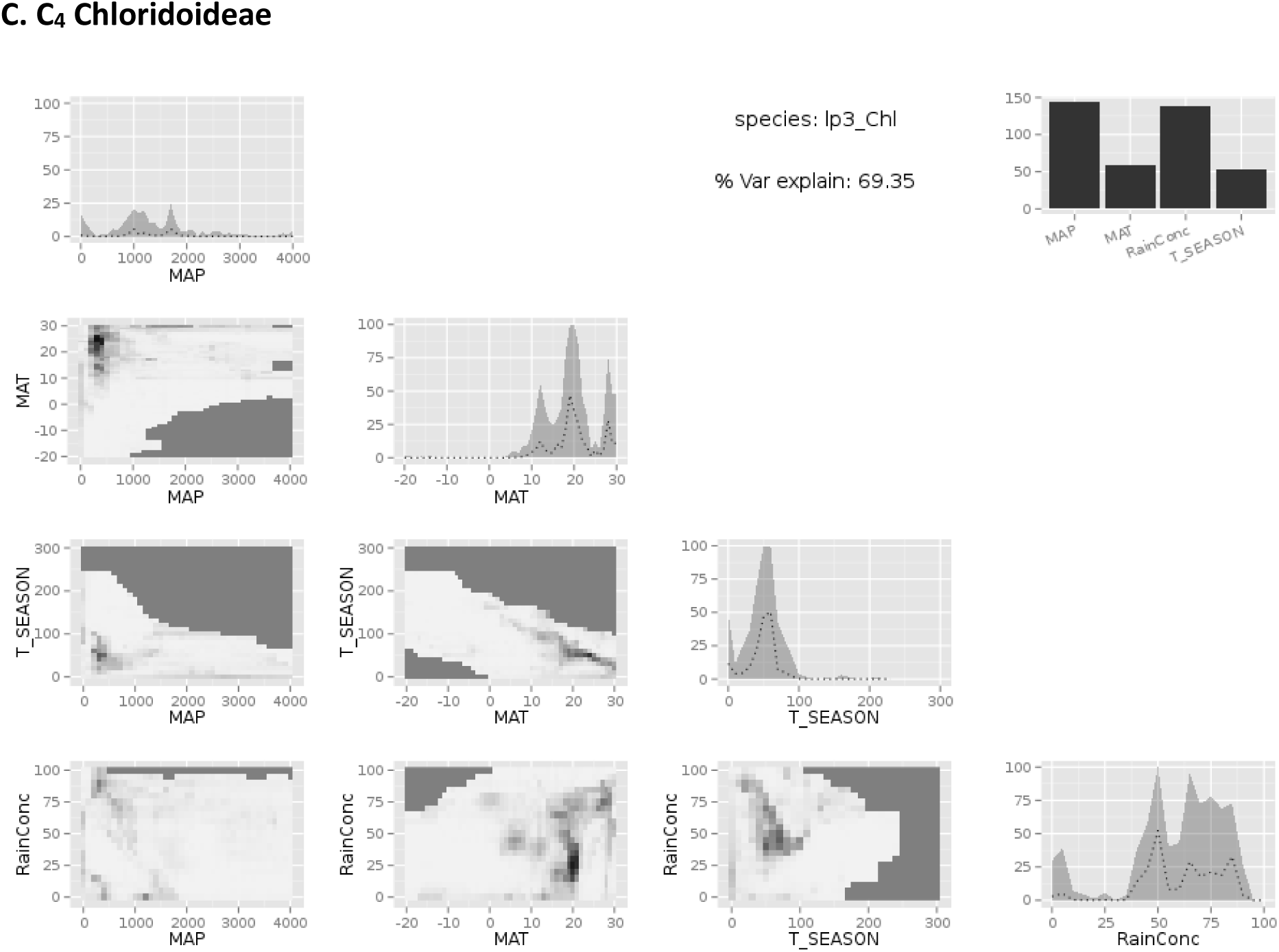
Plots from Random Forest analyses of the relative importance of mean temperature, mean precipitation, temperature seasonality and rainfall seasonality in the limits of the three key lineages of grasses. A) C3 BEP, B) C4 Andropogoneae, and C) C4 Chloridoideae. Model fits against data are shown for the land area over which each lineage dominates against each climate variable.

**Figure S6.**
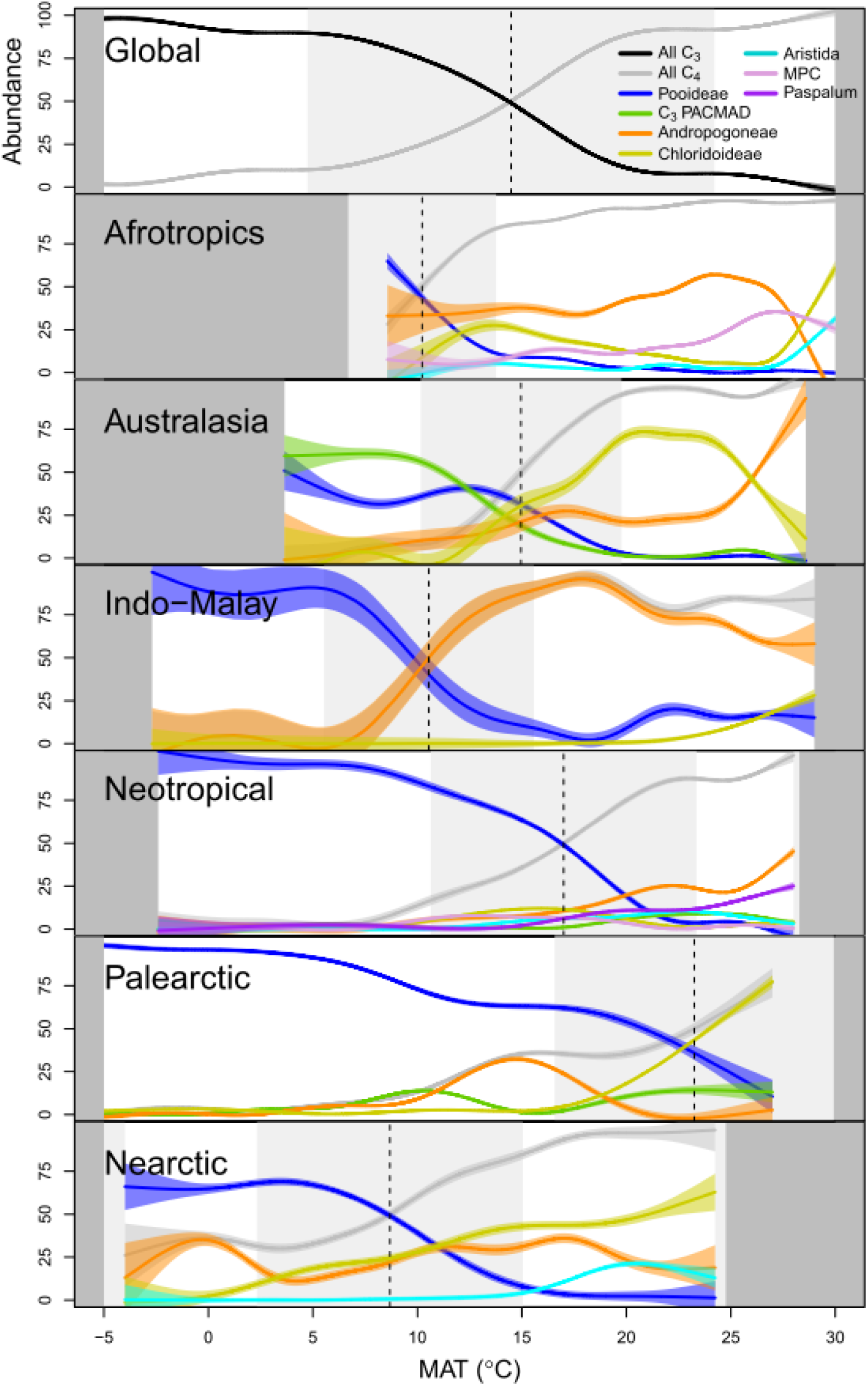
Lineage – temperature associations.

**Fig. S7:**
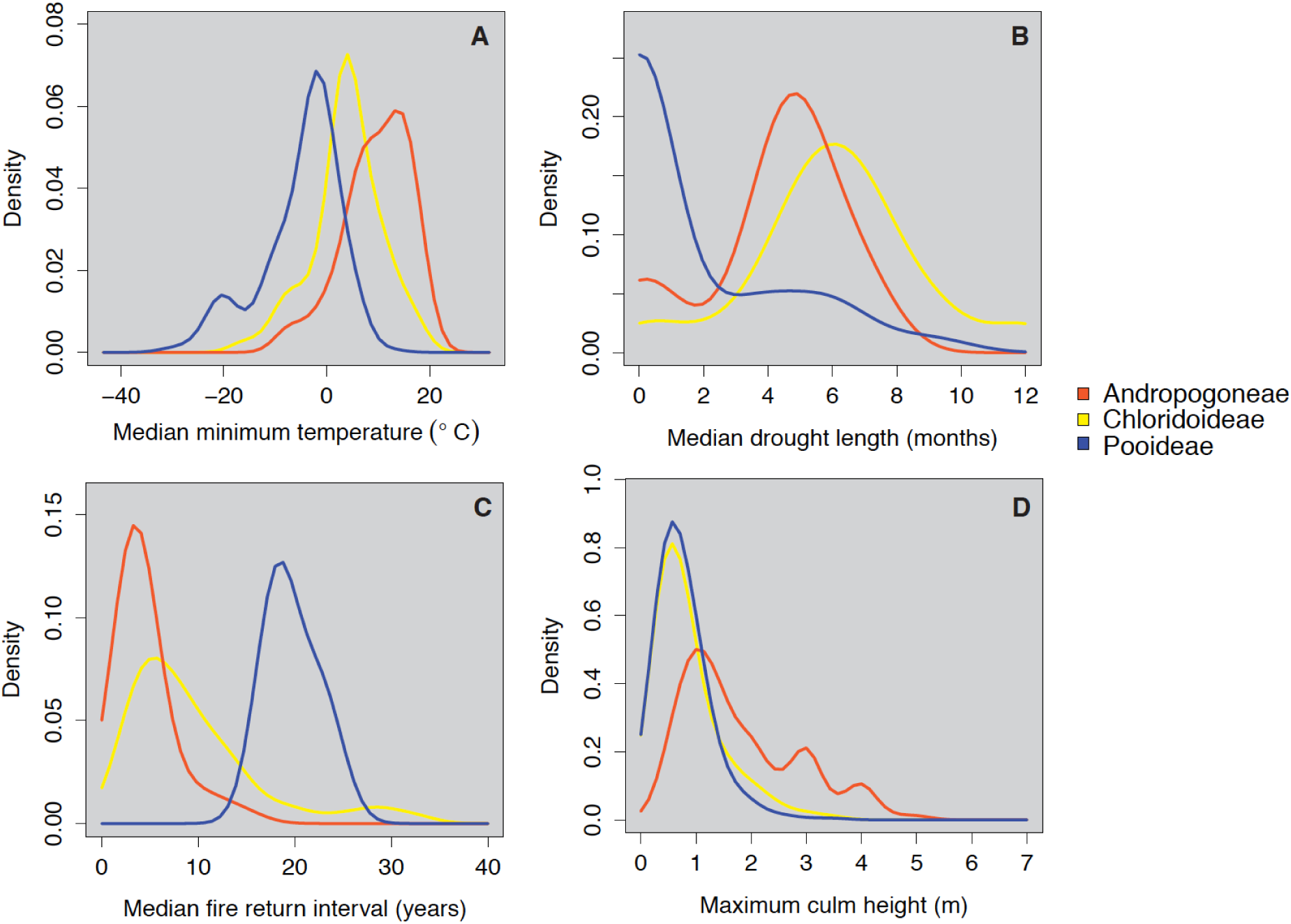
Trait-environment associations. Density plots of traits that characterise the realised ecological niche of dominant grass lineages: A) Median minimum annual temperature (°C) across the range of each species; B) Median drought length (months); and C) Median fire return interval (years), calculated by mapping GBIF occurrence data for each species onto Earth Observation data layers (see Methods). D) Maximum height of the culm (flowering stem) for each species, as a measure of plant size at maturity (see Methods). In each case, species from each lineage recorded within vegetation units in our dataset were mapped across their whole range (i.e. beyond the area over which they dominate ground cover).

**Figure S8.**
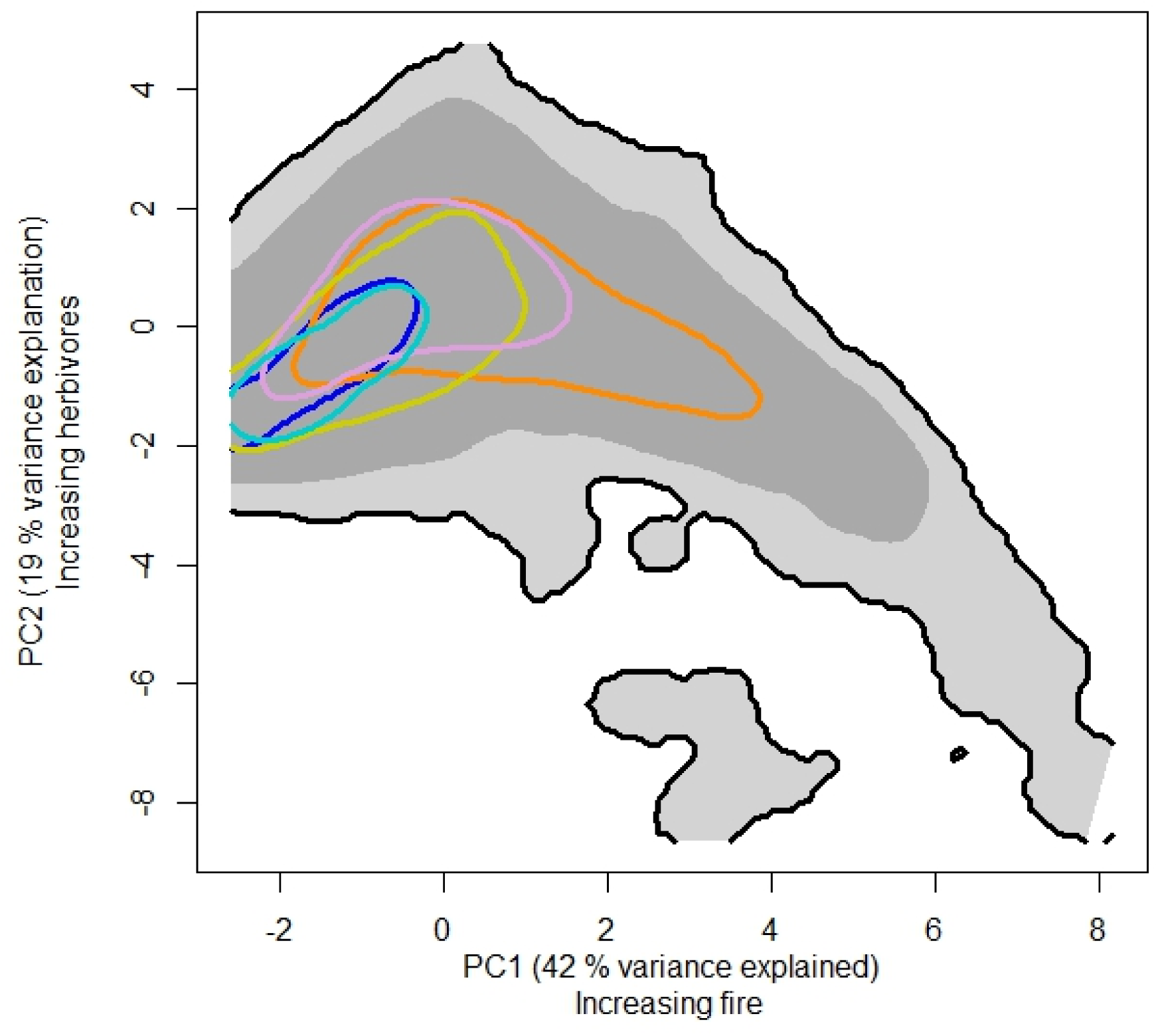
Ordination of fire frequency and herbivore biomass with the 90^th^ quantile of lineage distributions shown via different colours. This ordination shows the limits of five grass lineages relative to the prevalence of fire and modelled herbivore biomass. Colours representing lineages are consistent with figures in the main text. Key here is that one lineage stretches into environments of more frequent fire (Andropogoneae = orange), while a number of lineages are clustered and overlapping with respect to variation in herbivore biomass. However, it is worth noting the globally poor data on herbivore biomass in contrast to fire that is relatively easily quantified by satellites as changes in surface reflectance and heat.

**Figure S9.**
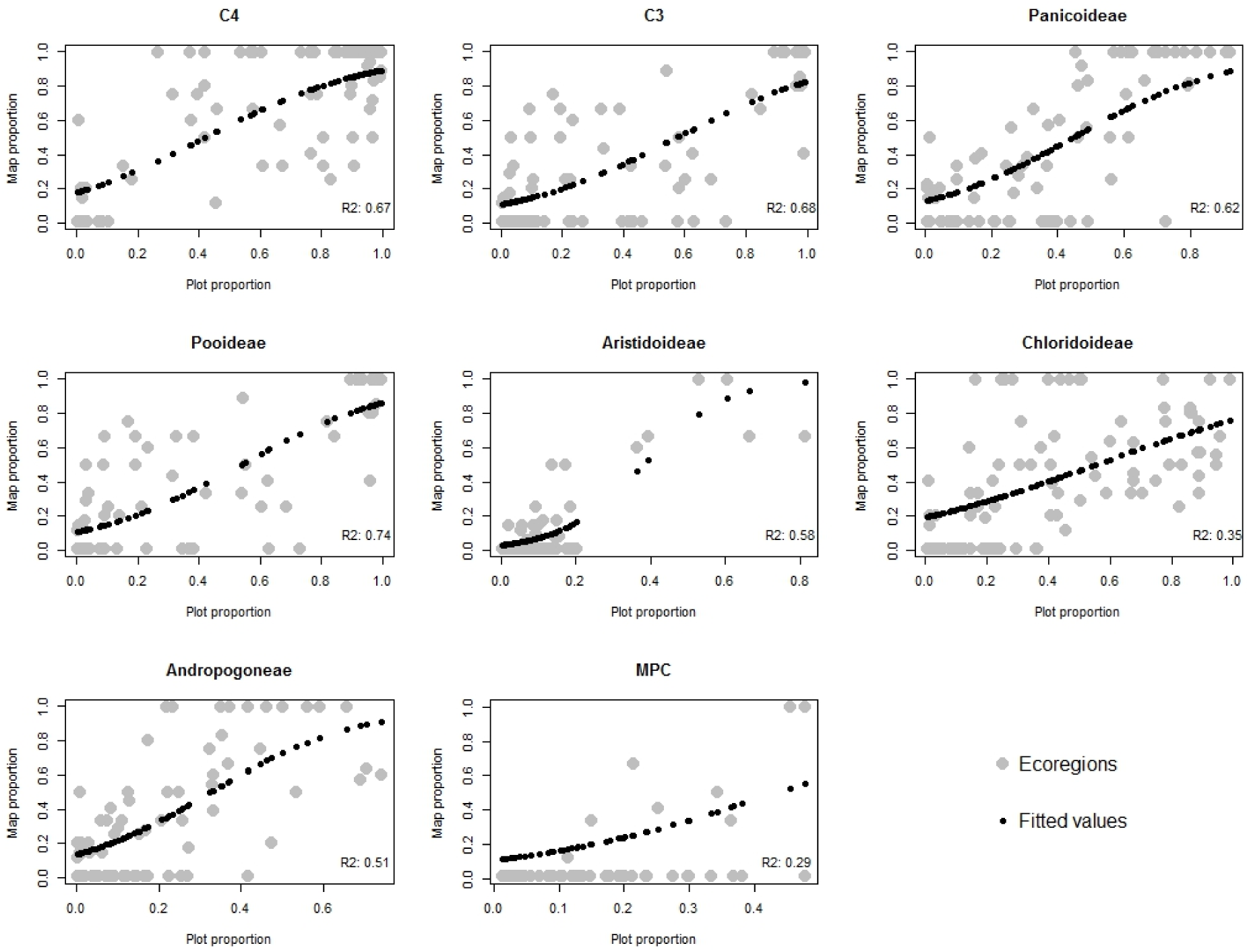
Validation of mapping approach to determine function and lineage level distributions of grassy biomes. Shown are figures relating plot level versus map level estimates of different grass groups (as shown on each plot). Logistic regression was used to quantify relationships and the deviance explained of the analyses are shown on each plot.

**Table S1:**
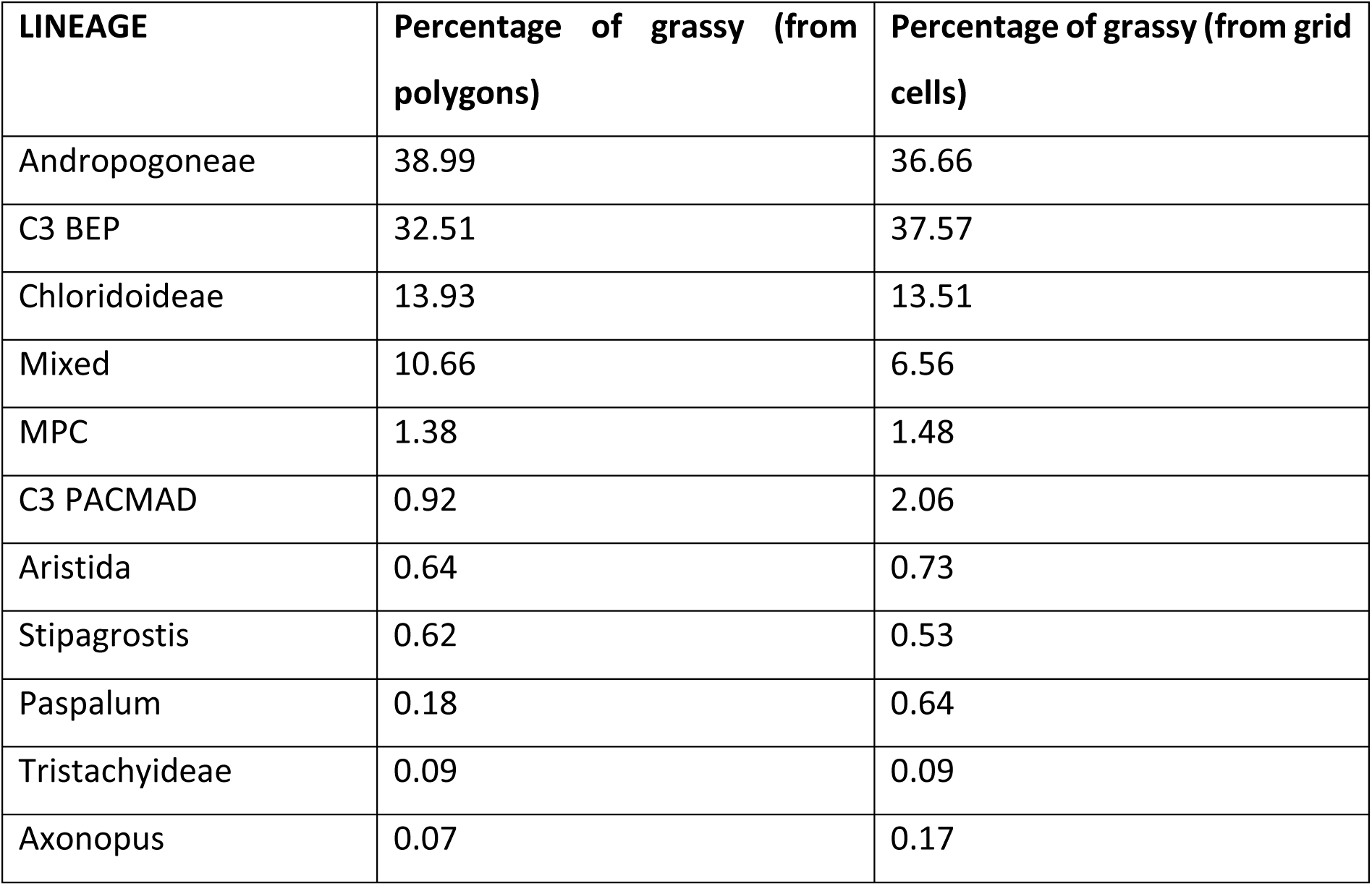
Land area occupied by each grass lineage. Each column represents a slightly different way to calculate the relative coverage of grassy biomes by different grass lineages. Polygon calculations are based on the mapped polygons while grid cells represent the conversion of data from Both calculations use a WGS84 projection.

